# A clustering neural network model of insect olfaction

**DOI:** 10.1101/226746

**Authors:** Cengiz Pehlevan, Alexander Genkin, Dmitri B. Chklovskii

**Affiliations:** Center for Computational Biology, Flatiron Institute, New York, NY; NYU Langone Medical Center, New York, NY

## Abstract

A key step in insect olfaction is the transformation of a dense representation of odors in a small population of neurons - projection neurons (PNs) of the antennal lobe - into a sparse representation in a much larger population of neurons -Kenyon cells (KCs) of the mushroom body. What computational purpose does this transformation serve? We propose that the PN-KC network implements an online clustering algorithm which we derive from the *k*-means cost function. The vector of PN-KC synaptic weights converging onto a given KC represents the corresponding cluster centroid. KC activities represent attribution indices, i.e. the degree to which a given odor presentation is attributed to each cluster. Remarkably, such clustering view of the PN-KC circuit naturally accounts for several of its salient features. First, attribution indices are nonnegative thus rationalizing rectification in KCs. Second, the constraint on the total sum of attribution indices for each presentation is enforced by a Lagrange multiplier identified with the activity of a single inhibitory interneuron reciprocally connected with KCs. Third, the soft-clustering version of our algorithm reproduces observed sparsity and overcompleteness of the KC representation which may optimize supervised classification downstream.

## I Introduction

In the quest to understand neural computation, olfaction presents an attractive target. Thanks to recent technological developments, there is a wealth of experimental data on olfactory processing. Stereotypy of early olfactory processing between vertebrates and invertebrates suggests the existence of universal principles. Yet, a comprehensive algorithmic theory of olfaction remains elusive.

Here, we attempt to understand the computational task solved by one key olfactory circuit in invertebrates [1], Figure 1, top: antennal lobe (AL) projection neurons (PNs) synapsing onto a much larger population of Kenyon cells (KCs) in the mushroom body (MB) which in turn are reciprocally inhibited by a single giant interneuron (GI, which is called GGN in the locust [2], [3] and APL in *Drosophila* [4], [5]). The PN-KC-GI circuit transforms a dense PN activity vector into a sparse representation in KCs [6], [7], [8]. The lack of a teaching signal suggests that this transformation is learned in an unsupervised setting. In contrast, learning the weights of KC synapses onto MB output neurons (MBONs) is thought to rely on reinforcement [9].

## II Related work

Many existing publications proposed mechanistic models of the PN-KC-GI circuit and attempted to explain known structural and functional features. In particular, they aimed to explain the sparseness of the KC activity [10], [2], [11], [12], [13] and studied how sparseness and other features affect olfactory learning and generalization at the output of KCs [14], [15], [16], [17]. A recent work proposed that KC representations solve a similarity search problem by representing similar odors with similar KC responses [18], following the similarity matching principle [19].

However, these papers did not attempt to derive the observed network structure and function from a principled computational objective. Such derivation is the goal of the present paper.

### A Demixing network models

Previous attempts to derive network structure and function assumed that the computational task of olfactory processing, in general [20], [21], [22], and the PN-KC-GI circuit, in particular [23], [24], [25], is compressive sensing or sparse autoencoding. In this view, odors are sparse vectors in the space of concentrations of all possible molecules. ORNs project these vectors onto the vectors of receptor affinities resulting in (normalized) vectors of AL PN activity. Then, the PN-KC projection recovers the concentration of each constituent molecule in the activity of a corresponding KC.

However, modeling olfaction with a compressive sensing or autoencoding computational objective has three problems. First, experimentally measured KC responses seem to contradict the demixing theory. Even though many KCs often respond selectively to a single component in an odor mixture [21], some KCs respond to specific mixtures but not to their components by themselves [21].

Second, circuits proposed for computing sparse representations contradict the PN-KC-GI circuit architecture. They require one of the following: an all-to-all KC connectivity [26], [27], [28], [29], many inhibitory interneurons [30], [31], [32], [33], [34], [35], or strictly feedforward architecture [23], [24], [20], all three inconsistent with the single inhibitory interneuron in the MB.

Third, demixing models often interpret nonnegative neural activity in KCs (spiking neurons cannot have negative firing rates) as a manifestation of the physical nonnegativity of molecular concentrations (counts of molecules cannot be negative) [36]. However, such explanation does not generalize to most other neurons in the brain whose activity is nevertheless nonnegative. Hence, there must be another reason.

**Fig. 1.**
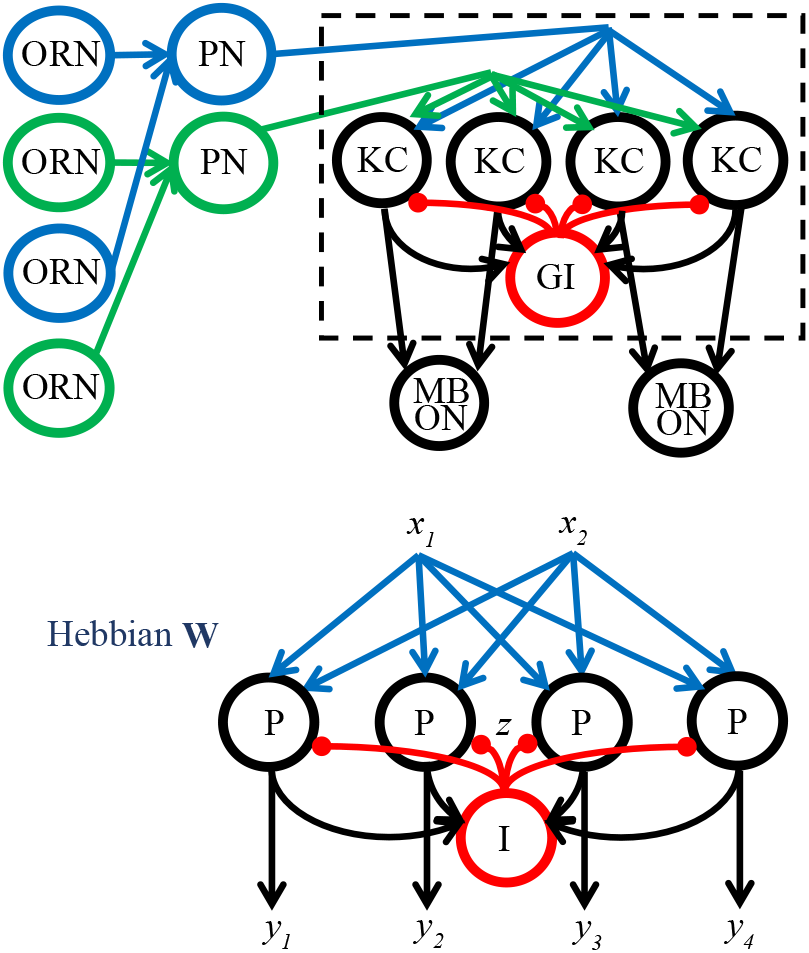
Top: Schematic of the insect olfactory system. Odors are transduced in the olfactory receptor neurons (ORNs). Each projection neurons (PN) receives inputs from ORNs of the same class in the AL glomerulus. Then, PNs (numbering ≈ 800 in the locust and ≈ 150 in *Drosophila*) synapse onto Kenyon cells (KCs) (numbering ≈ 50,000 in the locust [37] and ≈ 2,000 in *Drosophila* [9]) of the mushroom body (MB). A single giant interneuron (GI) provides reciprocal inhibition to KCs [4], [5]. KCs project onto a smaller number (34 in *Drosophila* [9]) of MB output neurons (MBONs). The dashed box delineates the circuit modeled in this paper. Bottom: A biologically plausible clustering network whose architecture is remarkably similar to the insect MB (P-principal neuron, I-interneuron).

### B Classic online *k*-means algorithm and neural networks

In this paper, we explore the hypothesis that the computational task of the PN-KC-GI circuit is to cluster streaming olfactory stimuli. In this subsection, we review a classic online clustering algorithm and clustering neural networks.

The goal of clustering is to segregate data into *k* classes. A popular clustering algorithm, *k*-means, [38], [39] achieves this goal by minimizing the sum of squared distances between data points, 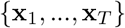, and cluster centroids, 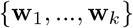, specified by the attribution indices 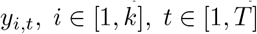:

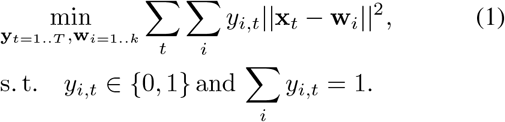

Here and below, vectors are lowercase boldface and matrices capital boldface.

Importantly, a biologically plausible algorithm must minimize Eq. (1) in the online (or streaming) setting. Indeed, sensory organs stream data to neural networks sequentially, one datum at a time, and the network has to compute the corresponding attribution indices on the fly without seeing the full dataset or storing any significant fraction of past inputs in memory.

The classic online *k*-means algorithm [38] performs alternating minimization of Eq. (1) by, first, assigning current datum, x_*t*_, to the cluster with the closest centroid, w_c_, i.e. “winner-take-all” (WTA), and, second, updating the number of data points in the winning cluster, n_c_, and the centroid of that cluster:

**Algorithm 1.**
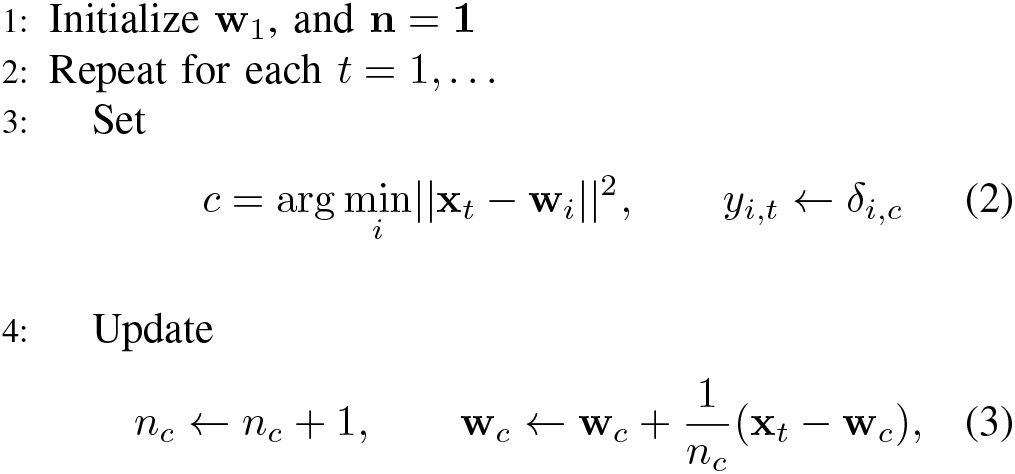
Classic online algorithm for *k*-means clustering

The online *k*-means algorithm is often given a neural interpretation [40], where *y_i,t_* is the activity of the *i^th^* output neuron (KC) in response to input x_*t*_ and w_*i*_ is the synaptic weight vector impinging onto that neuron. Yet, such interpretation is difficult to reconcile with known biology. First, computing squared distances in Eq. (2) seems to require the availability of x_*t*_ and w_*i*_ at the same location in a neuron. Contrary to that, while x_*t*_ is available only to the upstream neurons (PNs) and their synapses, the KC soma sees only 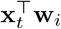. This difficulty could be circumvented by expanding the square and dropping the term, 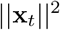, constant among all KCs. Then, the argument becomes 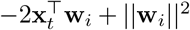 which could be computed in each KCs soma. Second, and more serious, difficulty is to actually find the minimum, (2), over all KCs. To compare 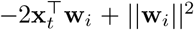 among KCs, the KCs would have to output that quantity, contradicting the assumption that they output *y_i,t_*.

Whereas neural implementations of clustering have been proposed before (reviewed in [40]), they either lack biological plausibility or are not derived from a principled objective. The central idea behind clustering neural network is competitive learning where neurons compete to assign a datum to their associated cluster by WTA dynamics. The “winner” neuron‘s synaptic weights, encoding that cluster’s centroid, are updated using a Hebbian rule. Examples of such algorithms include the classic *k*-means algorithm (see Section 2.3) [38], [39], the self-organizing maps (SOM) [41], the neural gas [42], the adaptive resonance theory (ART) [43], and the nonnegative similarity matching (NSM) network [28]. A probabilistic interpretation of WTA networks with Hebbian learning is given in [44].

## III Our contributions

This paper makes four main contributions: i) Starting from the classic *k*-means objective function, we derive an online algorithm that can be implemented by activity dynamics and synaptic plasticity rules in a biologically plausible neural network. In contrast, ART, SOM, the neural gas, and the classic *k*-means algorithms all assume that WTA operation is implemented in some biologically plausible way without specifying the network architecture or dynamics [45]. Hebbian plasticity, but not WTA dynamics, has been related to a clustering cost function in [46]. ii) The derived clustering network bears a striking resemblance to the architecture and dynamics of the PN-KC-GI circuit. Specifically, it accounts for the rectification in KCs as their activity represents nonnegative attribution indices and the existence of a single interneuron as its activity represents the Lagrange multiplier arising from the norm constraint (1). iii) In contrast with existing demixing models, we interpret activity of a KC as the presence of an olfactory object, which may be a complex odor made of many components or a simple odor of a single component, consistent with experimental findings [21]. iv) We extend our model to the soft-clustering scenario in which there could be multiple “winner” neurons. Such sparse, overcomplete representation is both reminiscent of the insect mushroom body and may be optimal for learning supervised classification downstream.

## IV A neural online *k*-means algorithm

In this Section, we present a new online *k*-means algorithm that overcomes both biological plausibility difficulties mentioned above.

### A Derivation of a neurally inspired online algorithm

We start by relaxing the constraint, 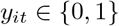, in (1):

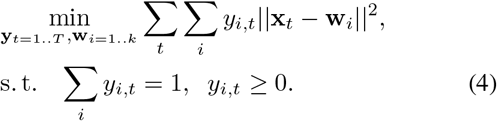

To see that, despite relaxation, the optimum of Eq. (4) has the property 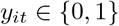, note that, for any value of w_i_s, the problem separates by data points. Then, for each *t*, we get:

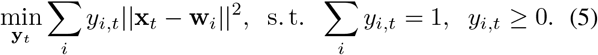

The solution of this problem is obviously, 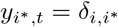 where 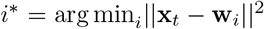, and 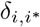 is the Kronecker delta, i.e., the same as for (1).

Next, we eliminate cluster centroids, w_i_s, from the objective. By taking a derivative of Eq. (4) with respect to wi and setting it to zero we find that the optimal cluster centroid is given by the center of mass, 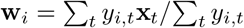. After substituting this expression into Eq. (4), some algebra, and using the Lagrangian form for the constraint, we obtain:

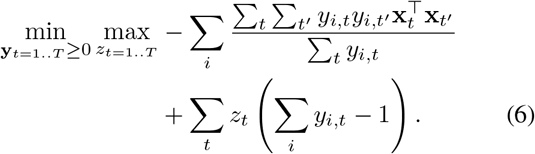

To derive an online algorithm, we reason that future inputs have not been seen and past outputs and Lagrange multipliers cannot be modified. Therefore, at each time step, *T*, we optimize Eq. (6) only with respect to the latest outputs 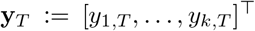 and the Lagrange multiplier, *z_T_*. Specifically, we solve:

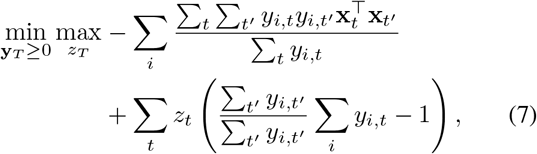

where we introduced the factor 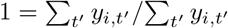 inside the parenthesis, which will be crucial for deriving an algorithm asymptotically equivalent to *k*-means (see Appendix I). Next, we drop i) the negligible current values from the sums in the denominators by invoking the large-*T* limit, ii) *y_i,T_* or *z_i,T_* independent terms, and iii) the negligible terms 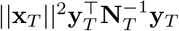 and 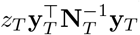 by again invoking the large-*T* limit. Finally, we arrive at:

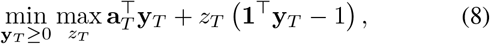

where

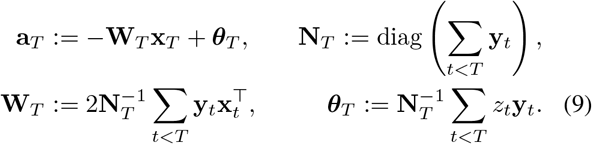

To guarantee invertibility of N*_T_* we initialize it with an identity matrix. We use a Matlab-inspired notation where, for any matrix M, diag(M) is a vector equal to M’s main diagonal and, for any vector v, diag(v) is a diagonal matrix with v on the main diagonal.

For each datum, our neurally-inspired Algorithm 2 performs two steps: i) Optimizes (8), ii) Recursively updates N*_T_*, W_*T*_ and *θ_T_* (Eq. (11) below) using optimal *y_T_* and *z_T_*. Because the optimal 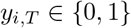, diag(N*_T_*) = n_*T*_ counts data points assigned to clusters until time, *T*.

**Algorithm 2.**
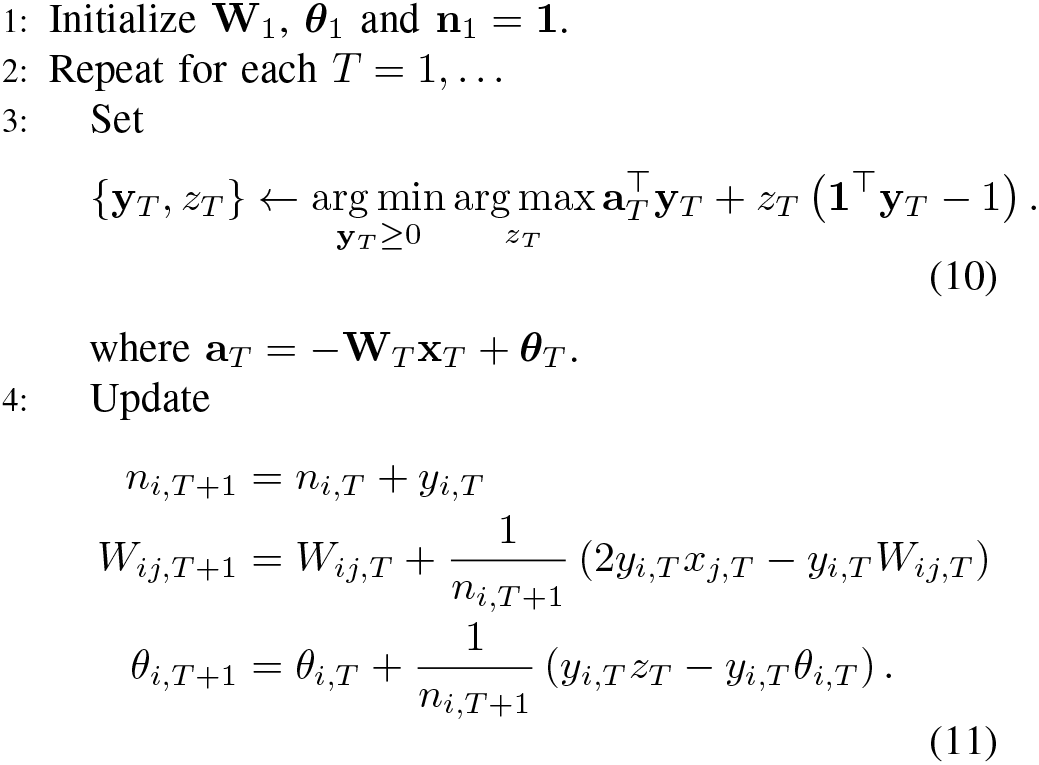
Neural algorithm for *k*-means clustering

Step 3 of Algorithm 2, i.e. Eq.(8), is the Lagrange form (with *z_T_* as the Lagrange multiplier) of the following optimization problem:

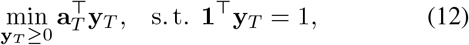

As Eq.(12) is a linear program on the simplex, it is optimized at the vertex with the smallest inner product value. Let *c* = arg min_*i*_ a_*iT*_, then 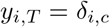, recovering the WTA rule.

To find the value of *z_T_*, we write the unconstrained Lagrangian: 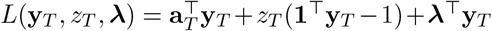. From complementary slackness, λ_c_ = 0 and, from stationarity, 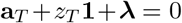 [47]. Therefore, 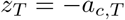 and we arrive at the following simplification:

Step 3 of Algorithm 2

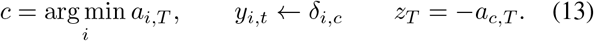

Comparing Algorithm 2 with classic *k*-means (Algorithm 1) we note several similarities. Cluster centroids (rows of ½W_*T*_) are centers of mass of the data points assigned to the corresponding clusters. Their update, (11), is identical to that in the classic online *k*-means algorithm (3). Both algorithms employ WTA dynamics.

Algorithm 2 differs from Algorithm 1 in that the “winner” is not determined by the Euclidean distance from cluster centroid to data point but by solving a linear program Eq. (12). Yet, despite this difference, Algorithm 2 is asymptotically equivalent to Algorithm 1 because 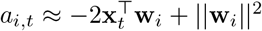 (see Appendix I). This solves the first difficulty with implementing *k*-means using neural networks.

Whereas Algorithm 2 with (13) gets us closer to a biological implementation by dropping the requirement for the availability of the terms 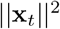 and 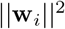 at each output neuron’s soma, it is still not clear how to implement the min operation biologically. The utility of Algorithm 2 with (13), which can be run on the CPU architecture, is due to its speed.

### B A neural circuit implementation

In order to implement the online algorithm 2 by a biologically plausible neural network, we solve the augmented Lagrangian of the linear program in Eq. (12) by primal-dual descent-ascent algorithm [47] (*ρ* is the augmented Lagrangian coefficient and *η* is the step size.):

Biologically plausible Step 3 of Algorithm 2

Iterate until convergence:

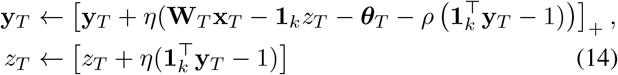

Using the dynamics in (14), Algorithm 2 has a biologically plausible neural network interpretation, Fig. 1, bottom. Each principal neuron (KC) receives input projected onto a synaptic weight vector (rows of W) representing the corresponding cluster centroid and the output activity of that neuron, *y_i,T_*, represents the corresponding cluster attribution index. *θ_i_*S are neural firing thresholds. The activity of a single inhibitory interneuron (GI) reciprocally connected with principal neurons represents the Lagrange multiplier, *z_T_*. The W update in Eq. (11) is a local Hebbian synaptic plasticity rule. The update of threshold, *θ*, is carried out by homeostatic plasticity. Finally, the augmented Lagrangian term may be implemented by the interaction via the common extracellular space shared by multiple principal neurons (KCs) [48], [49].

### C Numerical experiments

To evaluate the neural Algorithm 2 we compared it with the classic Algorithm 1 on the famous Fisher’s Iris dataset [50]. This dataset contains 150 samples equally representing three species of Iris: setosa, virginica, and versicolor. Each sample has four measurements: sepal length, sepal width, petals length, petal width (all in centimeters). We initialized the algorithms with three random points from a normal distribution with the mean and standard deviations of the whole data set, with independent coordinates. To emulate online behavior, the samples were permuted randomly. The same initial points and permutation were used for both algorithms.

The two algorithms perform similarly according to two indicators. Table I presents confusion matrices and Rand index values for the classic and neural algorithms, mean and standard deviation over 10 repetitions. Fig. 2, top, shows traces of cluster centroids evolution along the online execution of the two algorithms.

Fig. 2, bottom, gives an example of internal dynamics of the neural Algorithm 2, step 3 with Eq. (14), for one datum.

## V A neural soft *k*-means algorithm

Whereas our model explains the existence of a single inhibitory interneuron, it does not account for the observed activity of KCs. Indeed, in *k*-means clustering only one attribution index is non-zero for every datum implying that only one KC is active for each odor presentation. In contrast, experiments suggest that about 5-10% of KCs are active in the same time window [8].

### A Soft-clustering online algorithm and neural network

Our framework can explain this observation if, instead of hard clustering, we consider soft clustering where a datum may have multiple non-zero attribution indices. One may think that this problem can be solved by using the fuzzy *k*-means algorithm [52], where the cost function of Eq. (4) is generalized to

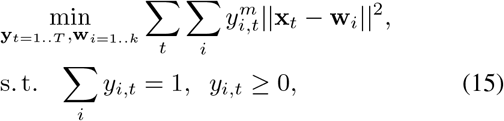

where *m* ≥ 1 (*m* = 1 limit recovers the hard *k*-means cost). However, fuzzy *k*-means does not produce sparse output, see Appendix II, in contradiction with experimental observations [6], [7], [8].

Therefore, we propose an alternative formulation which yields soft and sparse clustering and can be related to a semidefinite program (SDP) previously considered for clustering [53], [54] and manifold learning [55]. To this end, we add to the *k*-means cost function an l_2_-norm penalty:

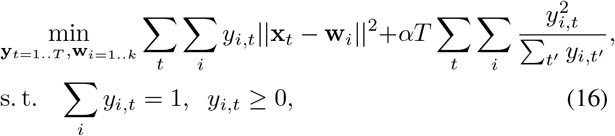

where the additional term in the objective is the only difference from (4), and *α* > 0 is a penalty parameter.

**TABLE 1.**
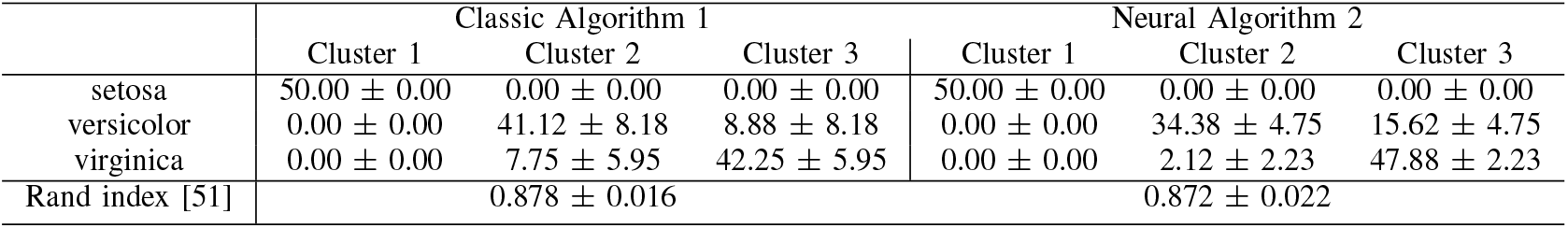
Confusion matrices on Fisher’s Iris dataset.

**Fig. 2.**
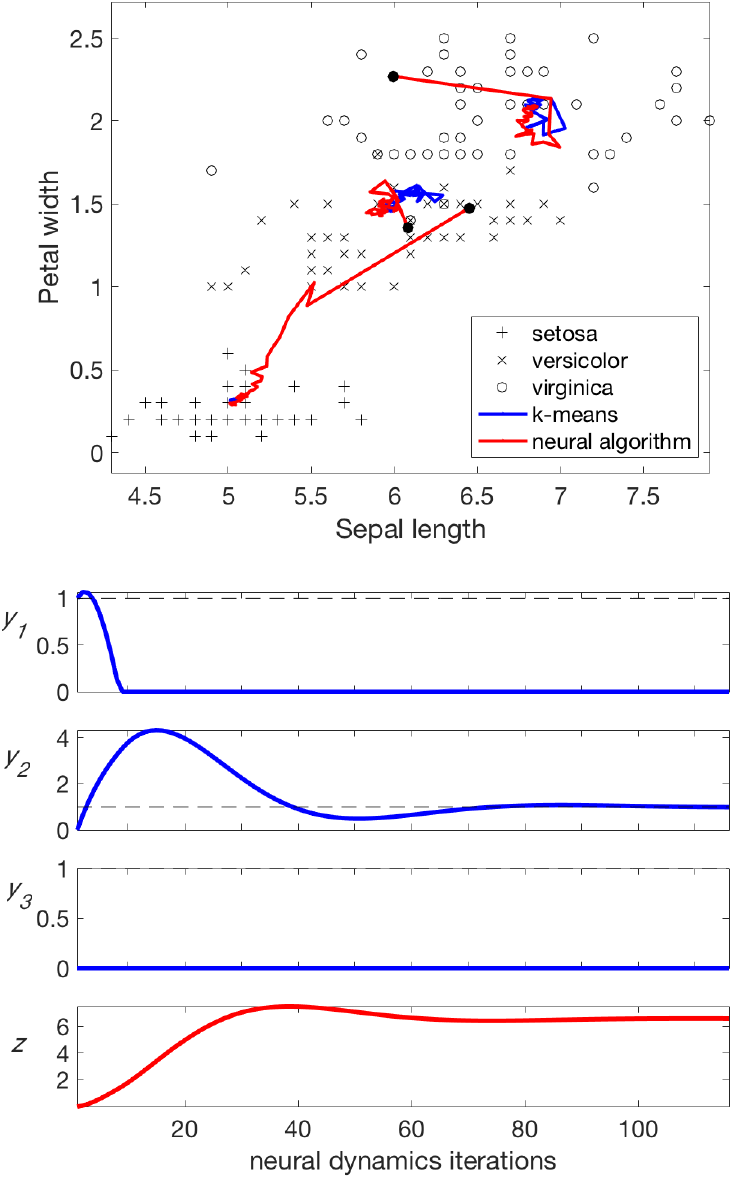
Performance of the neural Algorithm 2 on Fisher’s Iris data. Top: Trajectories of the three cluster centroids traced during the execution of the classic Algorithm 1 (blue) and the neural Algorithm 2 (red) initialized identically (solid black circles). In one case the traces almost coincide, in two other cases they deviate but end up in the similar positions. Shown is a projection of a 4D space onto 2D. Bottom: Neural dynamics during the execution of step 3 with Eq. (14), Algorithm 2, for one datum.

Following the steps in Section IV-A, we arrive at the online optimization problem for every time step:

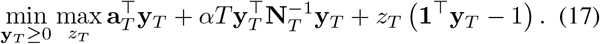

Solving optimization problem 17 and updating variables W, *θ* and N recursively, we arrive at the online neural Algorithm 3.

**Algorithm 3.**
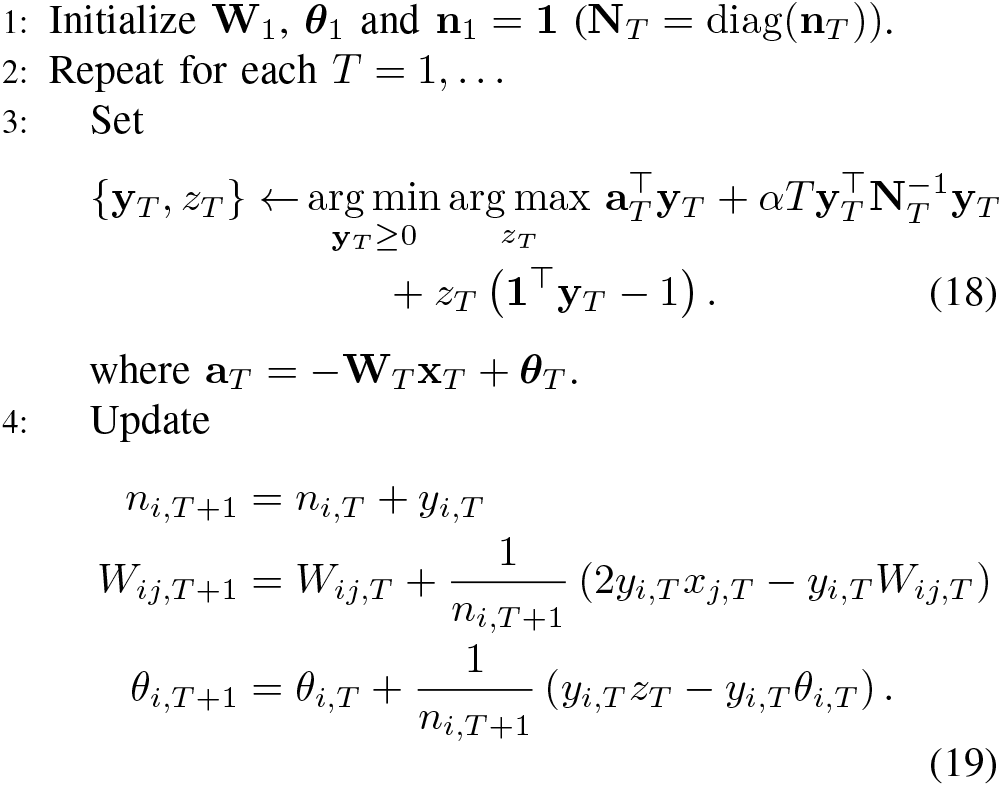
Neural algorithm for soft *k*-means clustering

Unlike Eq. (8), Eq. (17) is the Lagrangian form of a quadratic problem with linear constraints (with *z_T_* being the Lagrange multiplier):

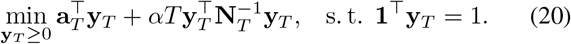

Step 3 of Algorithm 3 can be carried out by any quadratic program solver. To see why optimal solutions to (20) are sparse, note that for every *y_i,T_* > 0 the Lagrange multiplier for the non-negativity constraint is zero, so that the stationary condition takes the form:

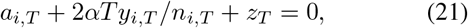

from which we have: 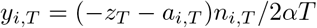. If all 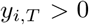, then using the equality constraint we can define: 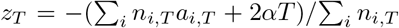. Using positivity of *y_i,T_* we have 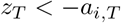, or

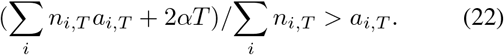

This inequality contains input variables only and cannot always be guaranteed to hold, so, in those cases, *y_i,T_* = 0 for some *i*.

In a biologically plausible neural circuit, the quadratic program, Eq. (20), in the augmented Lagrangian [47] form can be solved using primal-dual descent-ascent algorithm (*ρ -* the augmented Lagrangian coefficient and *η* - the step size):

Biologically plausible Step 3 of Algorithm 3

Iterate until convergence:

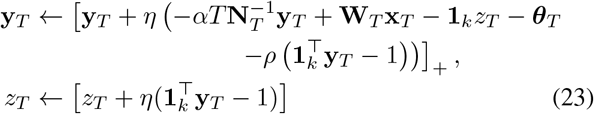

The only difference with the hard-clustering case, Eq.(14), is that the principal neurons are now leaky integrators with adaptive time constant, 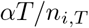.

### B Numerical experiments

We tested the utility of the representation learned by Algorithm 3 on the MNIST handwritten digits dataset by running a supervised classification algorithm on the vectors of attribution indices. The experiment was performed on a small-scale dataset (5,000 images were used in unsupervised training, last 1,000 images from unsupervised were used in supervised training, and another 1,000 images were used in testing) and on a large-scale dataset (60,000, 10,000, and 10,000 images, correspondingly). The results were averaged over 10 experiments with different random initializations.

After cluster centroids were learned by Algorithm 3 we classified the vectors of attribution indices using two versions of supervised learning that yielded similar results. The first was multiclass linear SVM, Figure 3. The second -directly connected each cluster to the most fitting target class (results not shown). Specifically, we defined *Td* as the set of all *t* values where the true digit is *d*. Then for each cluster *i* we assigned the best matching digit label by taking 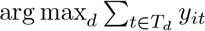. Then, we classified each image according to the digit which had the maximum sum of attribution indices.

We found that larger numbers of KCs led to better performance, Figure 3, top. Also, Figure 3, top, shows that nonzero values of the penalty parameter, α, resulted in better performance on small-scale (left) but not large-scale (right) datasets. Figure 3, bottom, shows that greater values of the penalty parameter, α, resulted in more active channels. This is consistent with viewing *α* as a regularization coefficient.

## VI Discussion

The proposed clustering neural network has a striking resemblance to the PN-KC-GI circuit in the insect olfactory system, Figure 1. We reproduced the general architecture including the divergence in the PN-KC circuit and, for the first time, derived the existence of a single inhibitory interneuron from a computational objective. The circuit dynamics in the model requires rectification in KCs, Eqs. (14,23), which occurs naturally in spiking neurons. The model does not require rectification in the inhibitory interneuron, (14,23), in agreement with the lack of spiking in giant interneurons [2].

According to our model, PN-KC synapses should obey Hebbian plasticity rules, Eqs. (11,19). KC-GI reciprocal synapses do not require plasticity, Eqs. (11,19). We predict interactions among KCs, Eqs. (14,23), that could be electrically mediated via shared extracellular space [48], [49].

**Fig. 3.**
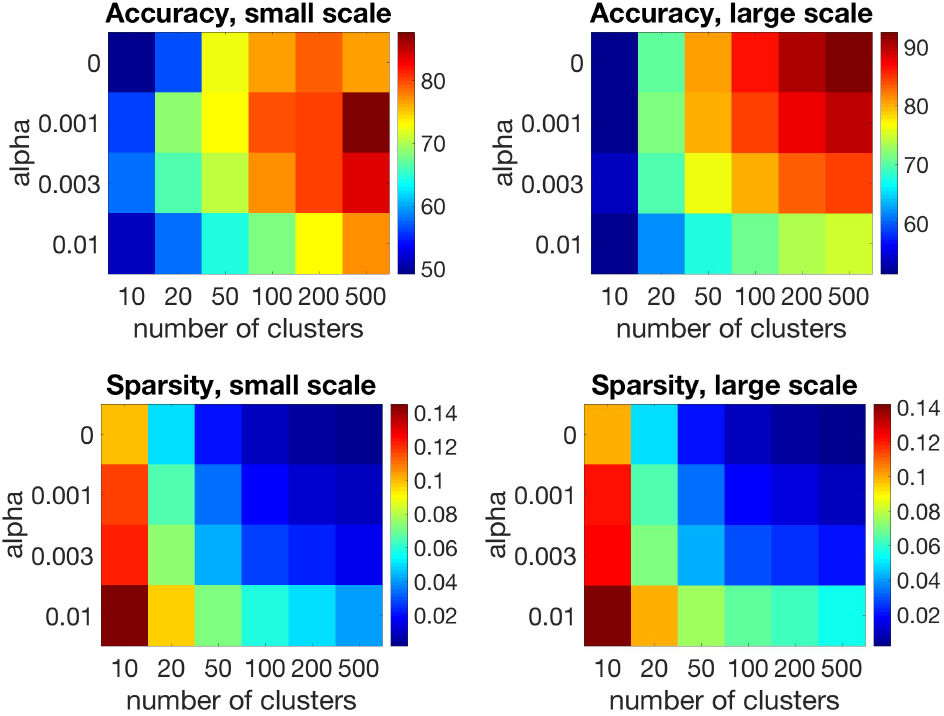
Performance of the soft clustering Algorithm 3 on the mnist hand-written digits data set for different *α* and number of clusters (special case, *α* = 0, corresponds to the hard-clustering Algorithm 2). Top: Accuracy of supervised digit classification based on attribution indices. Bottom: Sparseness of the attribution indices (average fraction of non-zero indices per datum).

In contrast to demixing models in which KCs signal the presence of constituent molecules in a complex odor, our model suggests that KCs represent olfactory objects which could be complex or simple odors. Our model predicts the existence of KCs selective to odor mixtures but not to the components alone. While preliminary evidence of such KC responses exists [21], they need to be investigated further.

Our clustering network has an advantage compared to existing random projection models [13], [17], [15], [18] in that it requires fewer KCs to represent olfactory stimuli. To span the PN activity vector space, random projections would require the number of KCs exponential in the number of dimensions. In contrast, our network is capable of reducing the required number of KCs by learning to represent only those parts of the PN activity space that are populated by data. In addition to minimizing volume [56] and energy [57] costs, reducing the number of KCs facilitates supervised learning.

In our model, nonnegativity of KC activity arises from the natural property of attribution indices (a datum cannot belong to a cluster with a negative attribution index). In contrast to the physical nonnegativity explanation of demixing models applicable only to neurons that code for molecular concentrations such as KCs, our interpretation can be applied, in addition, to most other spiking neurons encountered throughout the brain. Therefore, our interpretation is more general.

## Appendix I Neural clustering algorithm is asymptotically equivalent to classic *k*-means

Here, we demonstrate that Algorithm 2 asymptotically approaches classic *k*-means. To see this, let us focus on one cluster *i* and denote *t*_1_,*t*_2_, … the sequence of all time indices such that *y_i,t_j__* = 1. From Eq. (9) we have:

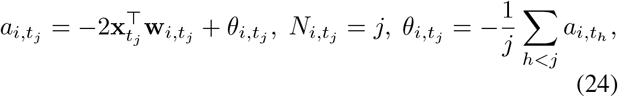

where we substituted the value of Lagrange multiplier as derived in the main text. Also, we denoted by w*_i,t_j__* the centroid of the *i*-th cluster, noting that the *i*-th row of matrix W*_t_j__* is double that vector. From the recursive relation between *a_i,t_j__* and *θ_i,t_j__* we can infer:

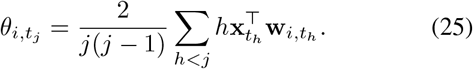

As the matrix, W, stabilizes with time, we make an approximation and replace the running cluster centroids with the latest value, and then take the expectation:

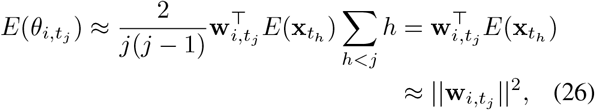

where for the last step we recall that vectors x_*th*_ are all and only those that constitute *i*-th cluster. Then

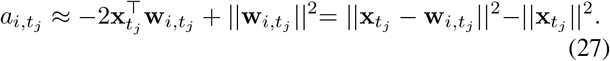

Therefore, asymptotically, the linear program solution of WTA in Algorithm 2 assigns each datum to the nearest cluster, just like *k*-means would, without requiring neurons to output Euclidean distances. Since the last term in (27) does not depend on *i*, the optimum in (6) only depends on distances between data points and, therefore, just like *k*-means, exhibits translational invariance (approximately), i.e. attribution indices depend only on the distance between data points, not on their absolute coordinates. In experiments, we observe the approximation to work well, so that *θ_i,t_j__* quickly approaches and, then, follows w*_i,t_j__* closely.

## Appendix II Fuzzy *k*-means produces dense output

To see why fuzzy *k*-means does not produce sparse output, write the Lagrangian for seeking *y_i,t_* under fixed w_*i*_:

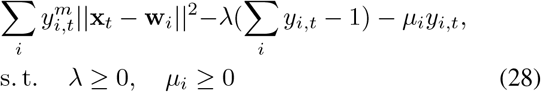

We write minus in front of λ because, since the objective is increasing, the values of *y_i,T_* need to be constrained only from below. Stationarity condition gives:

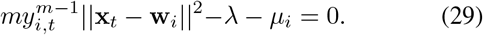

If 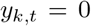, then stationarity becomes 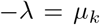, and due to non-negativity of both variables: 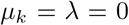. Then for all 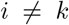 stationarity becomes 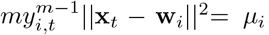, which, using complementary slackness, leads to all 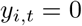 in violation of constraint unless for some *j* it happens that 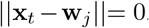. More generally, using some other than degree function *f*(*y_i,t_*) in the objective, a necessary condition for sparsity is *f*’(0) > 0.

## Appendix III Relation of *l_2_* -regularized *k*-means to an SDP for clustering and manifold learning

Here we show that the SDP used for clustering [53], [54] and manifold learning [55] is a convex relaxation of the *l_2_*-regularized *k*-means cost function (16).

To see this connection, we rewrite the optimization problem (16) in terms of new variables. Plugging in for optimal w_i_ = 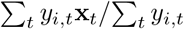, and defining new variables

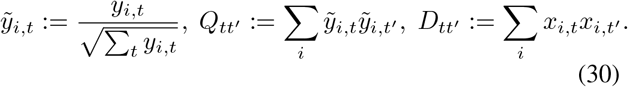

the cost (16) becomes 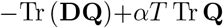. By construction, Q1 = 1 and Q ≥ 0. Therefore, we can relax the objective (16) to an SDP, where optimization is with respect to the positive semidefinite matrix Q:

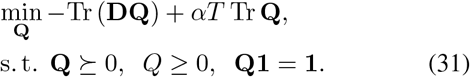

The SDP of [53], [54], [55] is

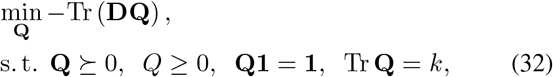

from which (31) can be recovered by replacing the last constraint by a penalty term in the cost.

## Acknowledgements

We thank Anirvan Sengupta, Mariano Tepper and Evgeniy Bauman for helpful discussions.

